# Enhanced Differentiation Potential of Pigmented Human Epidermal Equivalents

**DOI:** 10.1101/2025.05.29.656859

**Authors:** Yunlong Y. Jia, Julia Bajsert, Miguel Perez-Aso, Scott X. Atwood

## Abstract

Melanocyte-keratinocyte interactions are vital for regulating melanogenesis and maintaining epidermal homeostasis. However, most 3D human skin equivalents lack melanocytes, limiting their relevance for pigmentation studies. To address this, we utilized a pigmented human epidermal equivalent (PmtHEE) that incorporates melanocytes into the epidermis. Using single-cell RNA sequencing (scRNA-seq), we characterized PmtHEE and compared it with neonatal foreskin epidermis (FsEpi) and a fibroblast-containing human skin equivalent model (FibHSE). PmtHEE showed a higher proportion of differentiated cells and model-specific cell state transition paths that reflect *in vivo* trajectories. We further uncovered a shared L1CAM-EZR-driven external signaling network using exSigNet in FsEpi and PmtHEE that mediates melanocyte-to-keratinocyte communication and supports epidermal differentiation. Altogether, PmtHEE provides a distinct and physiologically relevant model of pigmented skin, with enhanced differentiation potential compared to conventional *in vitro* systems.

## Introduction

In human skin, interactions between melanocytes and keratinocytes are essential for regulating melanogenesis and melanin pigmentation. The epidermal melanin unit (EMU), consisting of a single melanocyte and approximately 36 of neighbouring keratinocytes ^1^, serves as a structural and functional collaborative unit within the epidermis. EMUs not only contribute to skin pigmentation but also plays a crucial role in maintaining epidermal barrier integrity ^2^, protecting against ultraviolet (UV)-induced damage ^3^, and modulating immune responses within the skin microenvironment ^4^. Dysregulation of pigmentary function has been implicated in various dermatological disorders, including melasma, vitiligo, and melanoma, highlighting the necessity for models that accurately recapitulate melanocyte behavior and pigmentation dynamics ^5,6^.

Contemporary three-dimensional (3D) human skin equivalent (HSE) models, primarily composed of fibroblasts (in the dermal layer) and keratinocytes (in the epidermal layer), are bioengineered *in vitro* tissues designed to mimic key aspects of skin structure and function ^7^. While these models offer advantages over traditional two-dimensional (2D) cultures, few incorporate additional supporting cell types such as melanocytes. Since the introduction of the first pigmented skin equivalent ^8^, substantial efforts have been made to refine and enhance *in vitro* pigmented skin models, resulting in significant advancements over the past four decades ^9–12^. To better replicate human skin pigmentation and its associated physiological functions, melanocyte-containing pigmented human epidermal equivalent (PmtHEE) models have gained traction due to their simplicity, reproducibility, and ability to incorporate melanocytes into the epidermal structure. The cellular behavior in PmtHEEs closely mirrors that of *in vivo* skin epidermis, with cells retaining the ability to respond to extrinsic regulatory stimuli, such as UV radiation ^13^.

PmtHEE models provide a promising platform for investigating pigmentation processes and the interplay between keratinocytes and melanocytes. However, conventional assessment methods, such as histological and immunofluorescence staining, offer limited insight into the molecular mechanisms and cellular heterogeneity of these systems. The advent of single-cell transcriptomics has revolutionized the skin field by enabling comprehensive, high-resolution analyses of cellular composition, states, and interactions ^14–16^. This approach has proven invaluable in characterizing organotypic culture systems and uncovering their strengths and limitations. Several studies have utilized single-cell omics to evaluate various skin organoids and organotypic models, yielding critical insights into their cellular and molecular characteristics ^17–19^. Despite these advances, no study has yet employed single-cell RNA sequencing (scRNA-seq) to characterize melanocyte-containing human skin *in vitro* models. Addressing this gap presents an opportunity to gain deeper insights into the molecular features of PmtHEE models and their fidelity to *in vivo* skin.

In this study, we successfully established a melanocyte-containing pigmented skin model that accurately replicates epidermal pigmentation *in vitro.* To further evaluate its resemblance to *in vivo* skin, we performed scRNA-seq analysis and conducted a comparative analysis with human neonatal foreskin epidermis (FsEpi) and a fibroblast-containing HSE model (FibHSE). Our findings demonstrate that PmtHEEs possess a distinct cellular composition and differentiation potential, underscoring their utility as robust platforms for studying pigmentation biology and advancing bioengineered skin substitutes.

## Results

### Characterisation of pigmented skin epidermal equivalents

A schematic overview of the culture process for the melanocyte-containing PmtHEE model is shown in **Figure 1A**. To initiate submerged culture, melanocytes and keratinocytes were mixed at a 1:20 ratio of melanocytes to keratinocytes and seeded onto an insert membrane. Following the formation of a confluent monolayer, removal of the apical medium established an air-liquid interface (ALI), triggering keratinocyte stratification. Although the PmtHEE model lacks a fibroblast-populated dermal layer, full epidermal formation is achieved within 14 days of ALI culture.

**Figure 1.**
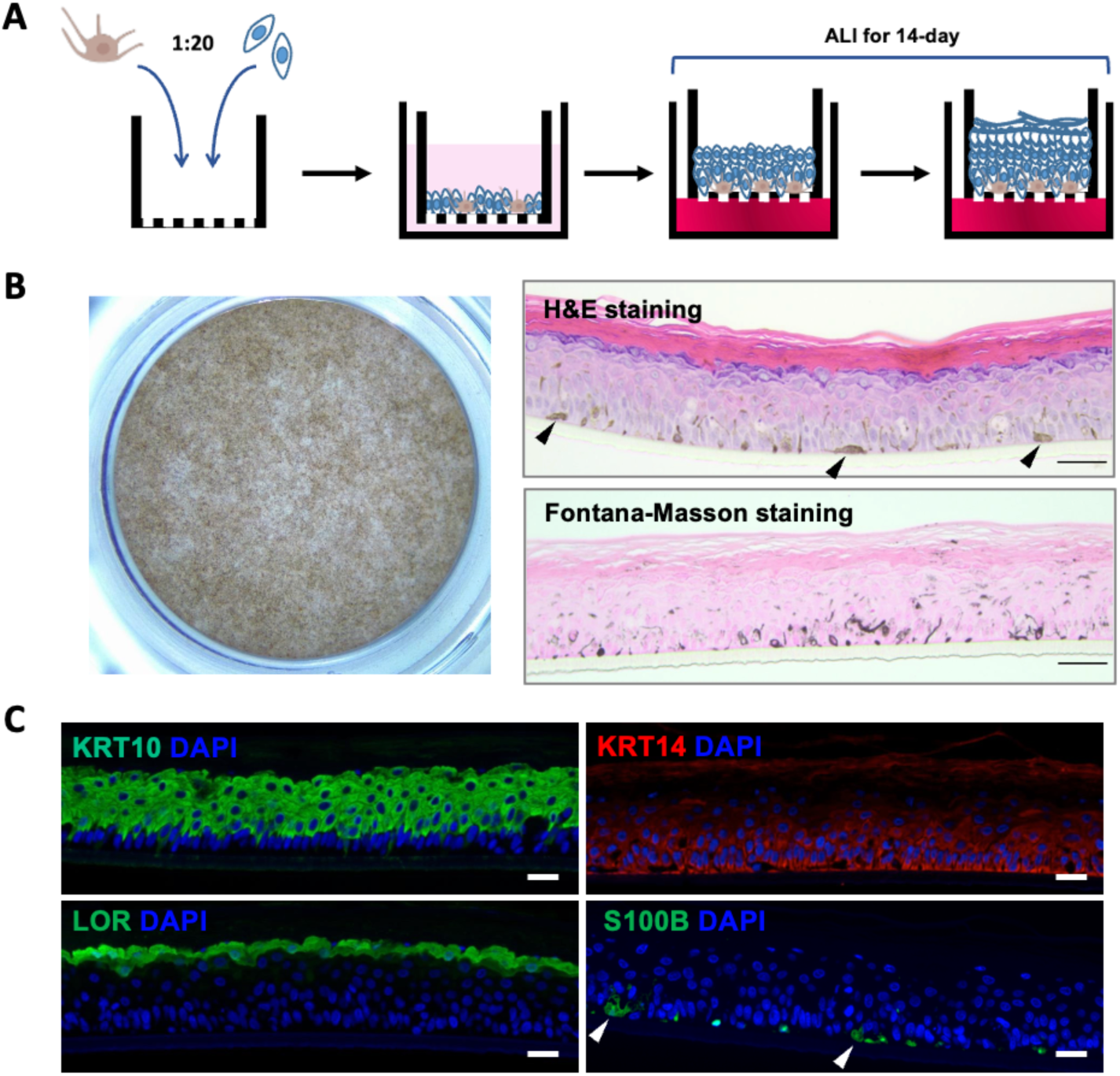
Generation and characterization of melanocyte-containing PmtHEE models. (A) Schematic representing the general formation of PmtHEEs. ALI, cultured at the air-liquid interface. (B) Melanin distribution in PmtHEEs. Top-view gross morphology of a PmtHEE after 14 days of ALI culture within the 24-well culture insert system (left), visualized using a stereomicroscope at 8x magnification. Histological analysis (right) includes haematoxylin and eosin (H&E) staining (top right), which illustrates epidermal morphology, and Fontana–Masson staining (bottom right), highlighting melanin distribution. Black arrowheads indicate regions of pigment accumulation. Scale bar, 50 µm. (C) Immunofluorescence analysis of KRT10, KRT14, LOR, and S100B expression. Nuclei are counterstained with DAPI (blue). White arrowheads indicate the positions of melanocytes. Scale bar, 50 µm.

A key feature of the mature pigmented skin model is its ability to more closely mimic the visual tone of human skin compared to classical melanocyte-free *in vitro* models. This pigmentation is visible to the naked eye and can be imaged using a stereomicroscope (**Figure 1B, left**). Histological analysis revealed a well-stratified and differentiated epidermis, with organized columnar keratinocytes in the stratum basale and a distinct stratum corneum (**Figure 1B, upper right**). Notably, pigmented cells were also observed within the stratum basale, resembling the distribution seen in native human skin. To better visualize melanocyte distribution and melanin dispersion, Fontana-Masson staining was performed (**Figure 1B, lower right**), revealing diffuse melanin deposition throughout the epidermis, indicative of melanin transfer to surrounding keratinocytes. Additionally, immunofluorescence (IF) analysis identified S100B-positive melanocytes localized within the basal layers of the epidermis (**Figure 1C**). To verify functional phenotypes and spatial localization of cell states within the model, we also analyzed key epidermal markers: KRT14 (basal), MKI67 (proliferation), KRT10 (suprabasal/spinous layers), and LOR, IVL, and FLG (granular/cornified layers) (**Figure 1C and S1**). The expression patterns were consistent with those observed in mature, healthy human skin.

Transmission electron microscopy (TEM) and ultrastructural analyses were undertaken to probe the architecture of the PmtHEE model in greater detail, focusing on the melanin deposition within adjacent keratinocytes. As shown in **Figure 2A**, the PmtHEE model exhibits morphological features that closely resemble those of native human skin. Notably, desmosomes, essential structures for maintaining stable intercellular cohesion ^20^, were observed at keratinocyte-keratinocyte junctions, and keratin bundles were localized around the perinuclear region. Regarding pigmentation, the detection of melanin within keratinocytes suggests active melanosome transfer and pigment dispersion from melanocytes, reflecting a physiologically relevant process. Consistent with observations in human skin ^21^, melanin organelles within keratinocytes were predominantly positioned in the perinuclear region (**Figure 2B**). As previously reported ^22^, only a limited number of melanocyte cell bodies are typically observed in ultrathin TEM sections of PmtHEE; however, dendritic extensions of melanocytes are frequently identifiable (**Figure 2C**). In line with the exo/phagocytosis model of constitutive pigmentation ^23^, where melanocores recently released from melanocyte dendrites were observed in close apposition to the plasma membrane of keratinocytes, suggesting ongoing pigment transfer.

**Figure 2.**
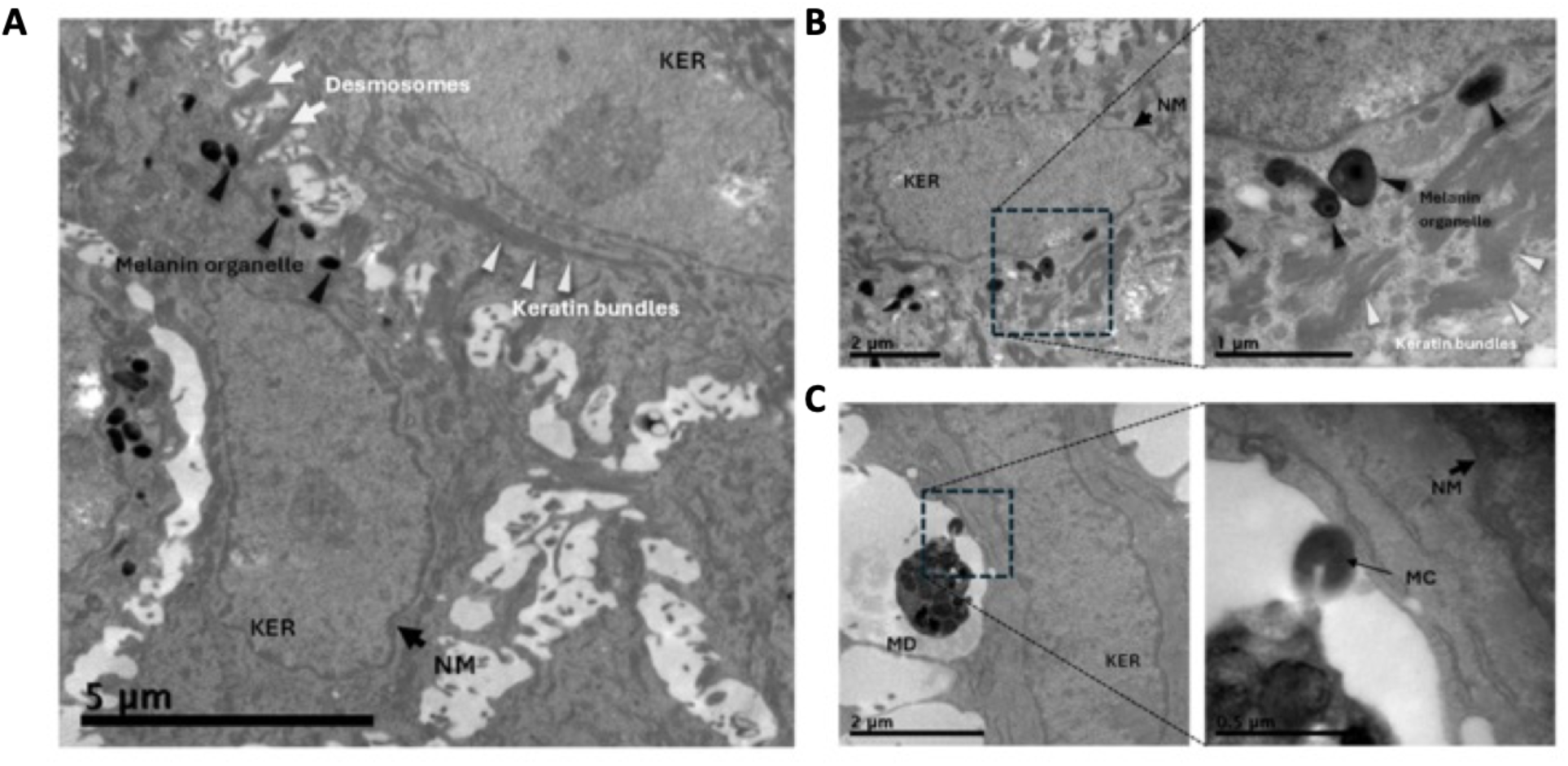
Ultrastructural characterization of the interface between melanin organelle and adjacent keratinocytes in the PmtHEE model. (A) Desmosomes (white arrows) connecting adjacent keratinocytes (KER). NM, nuclear membrane. Scale bar: 5 *µm.* (B) Melanin organelles are found in the perinuclear area of the KER, between the nuclear membrane (NM) and the keratin bundles (white arrowheads). Magnified picture of the boxed area is shown in the right panel. Scale bars: 2 *µm (left) and 1 µm (right).* (C) Melanocore (MC, thin black arrow) exocytosed from a melanocyte dendrite (MD) docking into KER’s plasma membrane. Magnified picture of the boxed area is shown in the right panel. Scale bars: 2 *µm (left) and 0.5 µm (right)*.

Collectively, these observations indicate that our PmtHEE model faithfully recapitulates the histological architecture and morphology of native human skin, exhibiting epidermal pigmentation comparable to that reported in previous studies ^22,24–26^.

### scRNA-seq reveals cellular heterogeneity and an enhanced suprabasal-like identity in PmtHEE models

We next performed scRNA-seq on our PmtHEE models and, in combination with previously published datasets ^19^, conducted a comparative analysis of single-cell transcriptomes from FsEpi, FibHSE, and PmtHEE models (**Figure S2**). Uniform Manifold Approximation and Projection (UMAP) visualization of the integrated Seurat object revealed six major cell types and multiple keratinocyte subpopulations (**Figure 3A**), including keratinocytes – comprising five basal states (BAS-I to BAS-V), four spinous states (SPN-I to SPN-IV), and three granular states (GRN-I to GRN-III) – as well as fibroblasts (Fib), melanocytes (MEL), hair follicle-associated cells (HF), Langerhans cells (LC), and endothelial cells (Endo). Cell type annotation was guided by canonical skin markers (**Figure 3C and S3**), as described in previous studies ^27–29^. Surprisingly, despite the intended melanocyte-to-keratinocyte ratio of 1:20 and the presence of melanocytes confirmed by histological analysis, only a single melanocyte was captured in the PmtHEE scRNA-seq dataset (**Figure 3B**).

**Figure 3.**
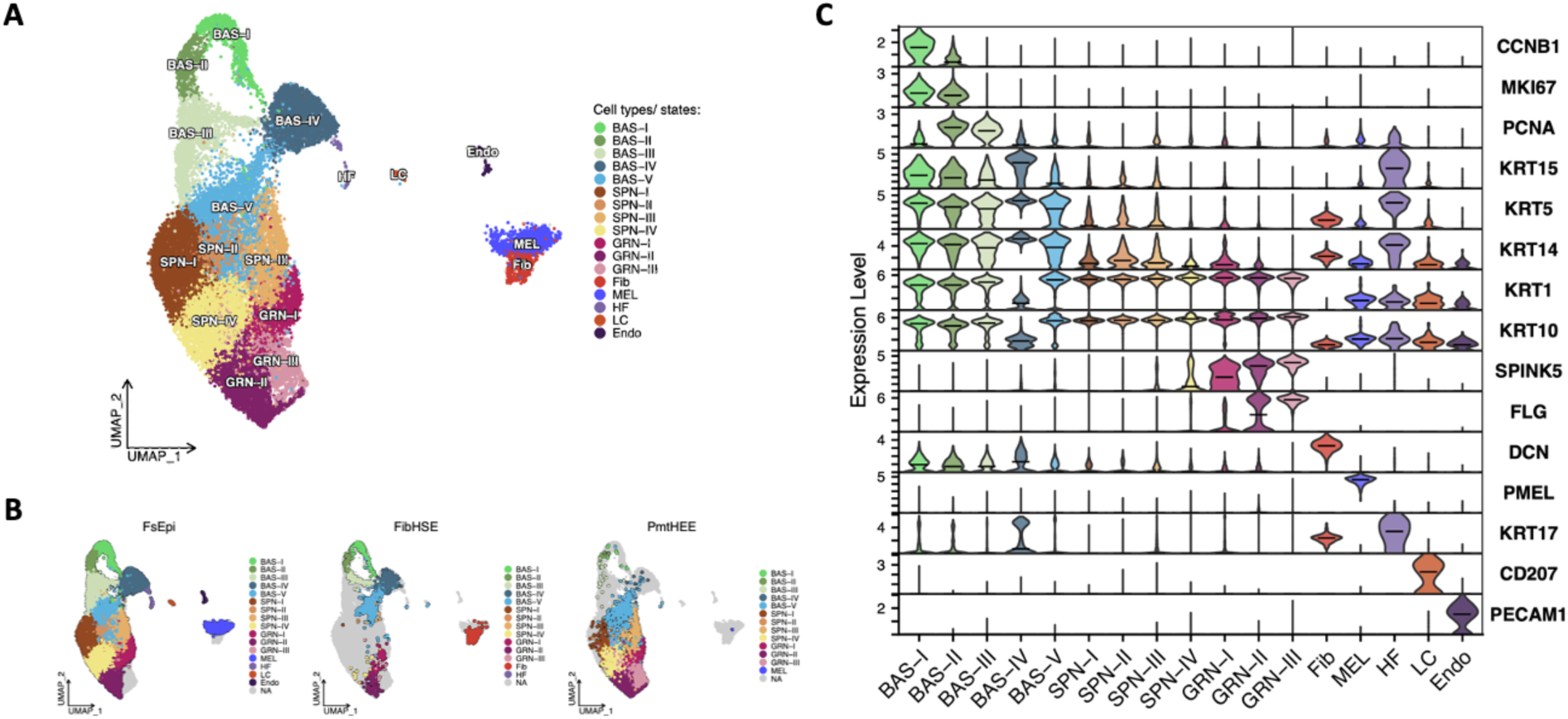
Annotation of cell populations in the integrated scRNA-seq dataset. (A) UMAP projection illustrating major cell types and identified keratinocyte subpopulations. (B) UMAP plots split by study, illustrating cell distribution across samples. (C) Violin plots showing the expression of predefined skin marker genes across identified cell populations. BAS, basal keratinocytes; SPN, spinous keratinocytes; GRN, granular keratinocytes; Fib, fibroblasts; MEL, melanocytes; HF, hair follicle-associated cells; LC, Langerhans cells; Endo, endothelial cells.

Although various bioengineered skin models have been developed to mimic the physiological complexity of human skin ^11^, keratinocyte-driven stratification and differentiation remain key determinants of overall *in vitro* skin morphology. Accordingly, a total of 21,106 keratinocytes were subsetted, re-clustered, and visualized using UMAP for downstream analysis (**Figure 4A**). To assess the similarity of keratinocyte states across different skin models, unsupervised hierarchical clustering was performed using MetaNeighbor ^30^, based on area under the receiver operating characteristic curve (AUROC) scores (**Data S1**). As shown in **Figure 4B**, the granular populations from PmtHEE, FibHSE, and FsEpi models exhibited high similarity. In contrast, PmtHEE-SPN displayed a lower degree of similarity when compared to the spinous populations of other models. Notably, PmtHEE-BAS demonstrated only weak similarity with basal keratinocytes from FsEpi and FibHSE, suggesting potential differences in basal state identity or heterogeneity. Cellular proportion analysis revealed an increase in spinous and granular keratinocytes, along with a decrease in basal cells in the PmtHEE model compared to the FibHSE model (**Figure 4C**). Consistent with the sparse distribution of BAS cells in PmtHEE, this may indicate that a subset of basal cells are undergoing state transitions and aligning more closely with other, more stable keratinocyte states rather than remaining within the basal compartment. Collectively, a stronger basal profile was observed in the FibHSE model, whereas PmtHEE exhibited a more suprabasal-like identity when compared to the balanced FsEpi control, as revealed by similarity scores and cell state composition analysis.

**Figure 4.**
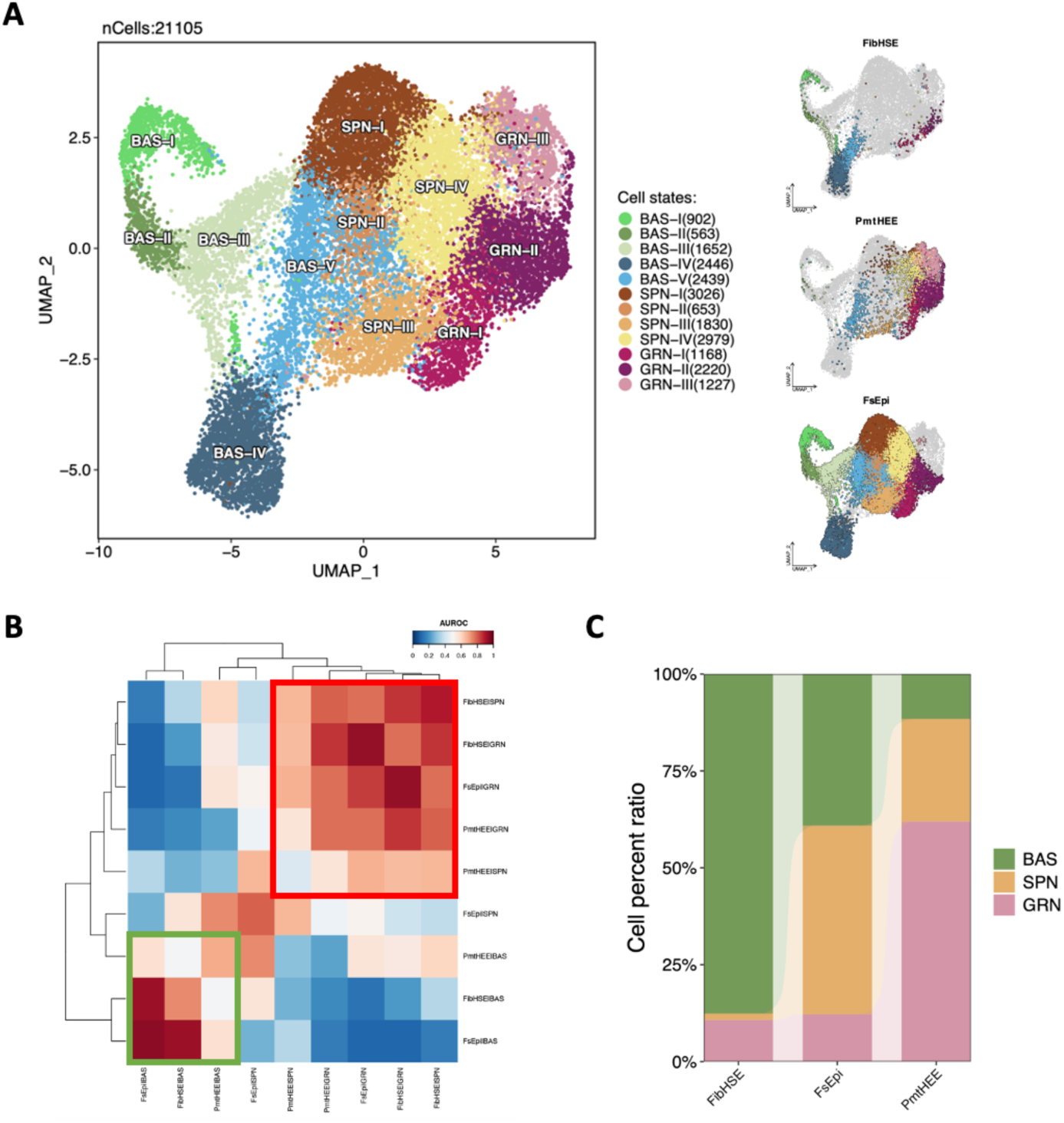
Characterization of keratinocyte cell states in the scRNA-seq dataset. (A) UMAP visualization of subsetted keratinocyte cell states. Contributions of each organoid to the overall UMAP is represented on the right. (B) Heatmap of MetaNeighbor analysis showing AUROC-based similarity across keratinocyte states. The red box highlights PmtHEE-associated suprabasal cell states, while the green box outlines PmtHEE-associated basal cell states. (C) Bar plot showing the proportion of keratinocyte cell states across different skin models. BAS, basal; SPN, spinous; GRN, granular.

### Shared branching trajectories in PmtHEE models are skewed toward suprabasal cell states

To investigate differences in keratinocyte phenotypic transitions across skin models, we performed pseudotime ordering and trajectory inference using Slingshot ^31^. In FsEpi, the highest number of branching events was observed, with five distinct lineages emerging from the BAS-II population. In contrast, FibHSE shared the same starting population (BAS-II) but exhibited only two predicted cell transition paths. Due to the underrepresentation of basal cells in PmtHEE, three lineages were inferred, originating instead from BAS-V (**Figure 5A and S4**). In summary, the combined results indicate that both FibHSE and PmtHEE partially share lineage trajectories with FsEpi (**Figure 5B**), exhibiting a basal lineage-enriched (BAS-II -> BAS-I -> BAS-III -> BAS-V) or suprabasal lineage-enriched profile (BAS-V -> SPN-II -> SPN-I; BAS-V -> SPN-III -> GRN-I; SPN-IV -> GRN-III), respectively. While the previously mentioned cell state imbalance may contribute to this lineage-specific bias, it also highlights the high fidelity of these *in vitro* skin models to their *in vivo* counterpart and their model-specific differences in cellular stemness potential.

**Figure 5.**
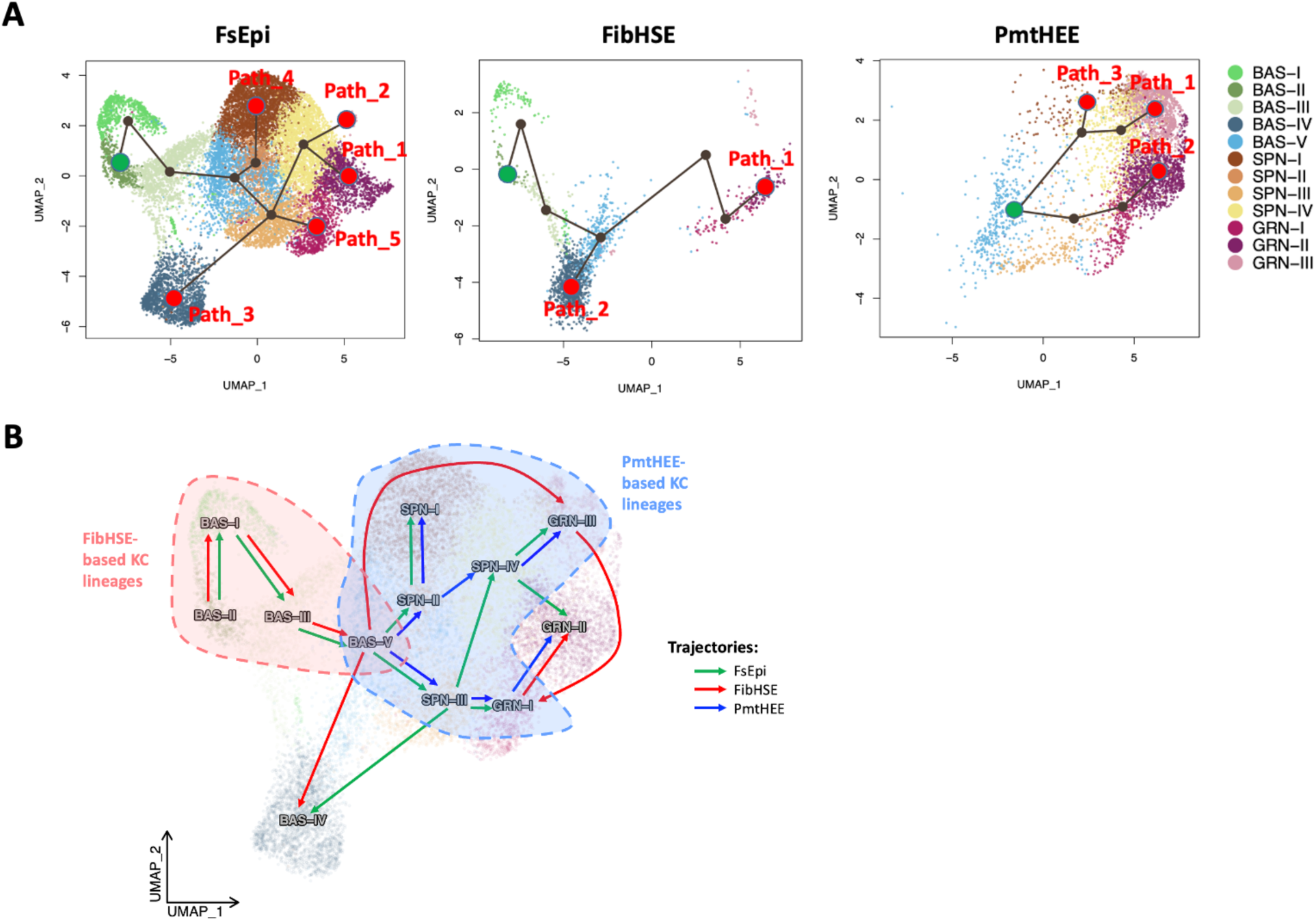
Slingshot-based inference of multiple branching lineages. (A) Pseudotime-based prediction of cell transition paths. (B) Comprehensive summary of lineage predictions based on identified keratinocyte (KC) states.

### The L1CAM-EZR-driven external signaling network mediates melanocyte-to-keratinocyte interaction in PmtHEE

Different skin models contain distinct cell types (**Figure 6A**), which underlie differences in cell-cell communication (CCC) and form the foundation for their versatility in various clinical and dermatological research applications ^32^. For example, FsEpi, serving as an *in vivo* reference model, primarily consists of keratinocytes, melanocytes, and fibroblasts. Though fibroblasts are absent in our dataset, downstream ligand-target interactions are still inferable ^33^. In contrast, FibHSE and PmtHEE, as *in vitro* models, each incorporate keratinocytes in combination with fibroblasts or melanocytes, respectively. This dual-component design not only reconstructs distinct morphological features but also drives model-specific cellular interactions, which in turn influence intra-model homeostasis – particularly in terms of keratinocyte dynamics. Thus, we hypothesize that PmtHEE faithfully recapitulates melanocyte-to-keratinocyte interactions.

**Figure 6.**
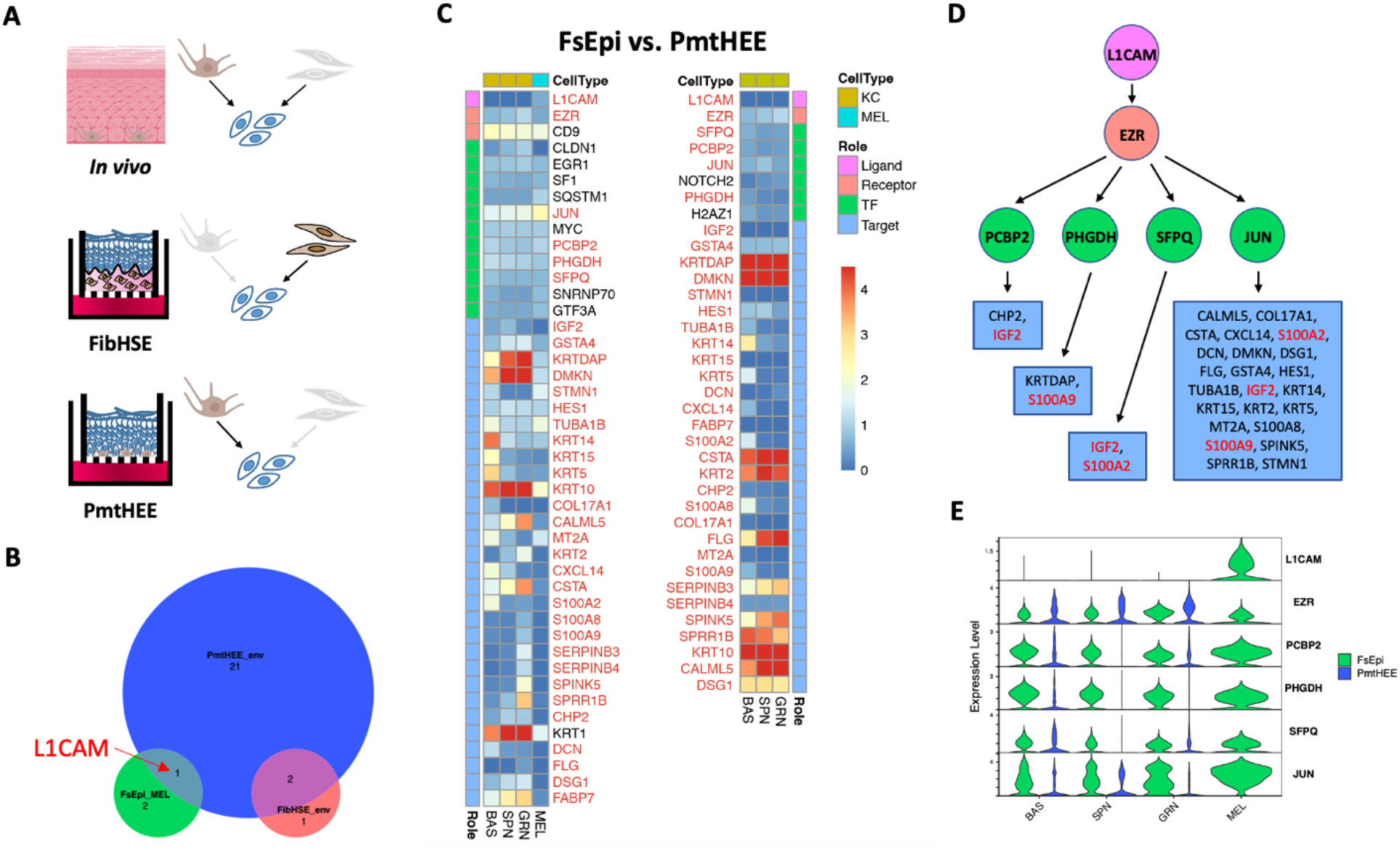
Melanocyte-to-keratinocyte interaction identified in FsEpi and PmtHEE conditions. (A) Schematic illustration of cell-cell communication across different skin models. (B) Venn diagram showing the predicted ligands under different culture conditions, based on exFINDER analysis. ’FsEpi_MEL’ represents ligands derived from melanocytes in *in vivo* skin; ’PmtHEE_env’ refers to ligands from the external environment (including melanocytes) in the PmtHEE model; and ’FibHSE_env’ denotes ligands from the external environment in the FibHSE culture model. Keratinocytes were used as the target cells in all analyses. (C) Heatmap of the expression levels of components in the L1CAM-EZR-driven exSigNet under FsEpi and PmtHEE conditions. Shared components are highlighted in red. (D) Topology plot showing the shared L1CAM-EZR-driven exSigNet, constructed using only shared genes. Repeated target genes were marked in red. (E) Violin plot showing the expression of ligand-receptor-transcription factor components from the shared L1CAM-EZR-driven exSigNet.

Using exFINDER ^33^, we defined keratinocytes – including BAS, SPN, and GRN states – as target cells to infer melanocyte-specific external signaling networks (exSigNets) across different models (**Figure S5A**). Among all ligands, L1CAM was the only melanocyte-derived ligand shared between FsEpi and PmtHEE (**Figure 6B**). The components of the L1CAM-driven exSigNet were further visualized to illustrate differences in downstream regulatory elements (**Figure 6C**). Notably, a single L1CAM-EZR ligand-receptor pair was identified that recapitulated *in vivo* interactions. This axis connects L1CAM on melanocytes with EZR-associated signaling platforms in keratinocytes and activates a conserved transcriptional network involving four TFs: PCBP2, PHGDH, SFPQ, and JUN. Nearly all downstream target genes were shared across models, suggesting a robust and conserved communication route. Consistent with previous reports on L1CAM-EZR interaction ^34^, this pathway may function through NF-κB activation, a signaling cascade known to regulate the balance between keratinocyte proliferation and differentiation ^35^. Within the shared L1CAM-EZR-driven exSigNet (**Figure 6D**), JUN emerged as the dominant transcription factor, regulating the majority of target genes. As a core component of the AP-1 transcription factor complex, JUN governs genes involved in proliferation, differentiation, and apoptosis – processes critical to skin development and renewal ^36^. While PHGDH, PCBP2, and SFPQ contribute to metabolic and stress-responsive pathways ^37–39^, they do not substitute for the master regulatory role of JUN in skin-specific transcriptional programs. Based on gene expression comparisons between FsEpi and PmtHEE (**Figure 6E**), JUN was the only transcription factor consistently expressed across all keratinocyte cell states. Its broad expression may facilitate the BAS-to-SPN/GRN transition, potentially contributing to the observed shift in cellular composition described earlier.

Taken together, although direct biochemical evidence for L1CAM-EZR binding in skin is limited, computational predictions and CCC analyses from scRNA-seq data strongly support its potential involvement in a broader signaling network that mediates melanocyte-to-keratinocyte paracrine communication and may regulate epidermal differentiation (**Figure S5B**) – possibly contributing to the enhanced differentiation potential observed in the PmtHEE model.

## Discussion

Organotypic skin culture models have been the most widely used *in vitro* 3D skin systems since their initial development in the 1980s ^40,41^. Evolving from simple keratinocyte-only human epidermal equivalents to more complex HSEs incorporating multiple cell types, substantial efforts have been made to replicate various aspects of skin physiobiology and morphology for both research and clinical applications ^42–44^.

Compared to traditional histological assessments, scRNA-seq offers an unbiased, high-resolution approach for benchmarking *in vitro* skin models. It enables comprehensive characterization of cellular composition, lineage dynamics, and molecular fidelity relative to native human skin. The emergence of cyst-like, hair-bearing skin organoids derived from human pluripotent stem cells has gained considerable attention ^18,45^. Through integration with single-cell transcriptomics, these organoids demonstrate the potential to function as a nearly complete, self-organizing *in vitro* skin system that faithfully recapitulates the hierarchical architecture of native skin – including stratified epidermis, adipocyte-rich dermis, pigmented hair follicles, and sebaceous glands. This advancement prompts a timely and important question: beyond the imperative to incorporate single-cell omics into current skin bioengineering workflows, how well do conventional models perform when evaluated at molecular and cellular resolution? And are they still sufficient to meet the increasingly complex demands of modern skin biology and translational research?

Within this context, our previous study profiled the single-cell landscape of various FibHSEs ^19^. Even in suboptimal models, we observed that cellular and signaling heterogeneity remained largely conserved and comparable to *in vivo* skin, underscoring their potential for modeling both general skin morphology and molecular perturbations. In the present study, we evaluated the melanocyte-containing PmtHEE model at both histological and single-cell levels. From a macroscopic perspective, PmtHEE exhibited a well-stratified and pigmented epidermis, characterized by low structural complexity but high molecular fidelity. Notably, it emerged as a highly reproducible model, surpassing previously reported FibHSEs in consistency and performance.

The diverse cellular compositions across the three models pose challenges for computational inference of CCC. Although only a single melanocyte was detected in the PmtHEE single-cell dataset, CellChat ^32^ remains compatible with cell groups containing equal or more than one cell, allowing for limited interpretation. Overall, FsHSEs exhibited a greater number of activated CCCs than PmtHEEs (**Figure S6**), as reflected in both the total number of interactions and the predicted signaling flow. Notably, pathways such as ncWNT, SLURP, and CXCL – dominantly active in PmtHEE – are closely associated with epidermal differentiation, immune modulation, and barrier function ^46–48^. In contrast, FsEpi displays a more diverse profile involving multiple classical signaling families, including those regulating proliferation and regeneration (e.g.,PTN ^49^), barrier maintenance and desquamation (KLK^50^), innate immune responses (e.g., TWEAK^51^), and vascularization and pigmentation (EDN^52^). These findings further support the enhanced differentiation capacity of the PmtHEE model.

However, a major limitation of CellChat analysis in PmtHEE lies in the low confidence of melanocyte-related signaling predictions due to the absence of a clearly defined melanocyte cell group. The underrepresentation of melanocytes and their limited transcriptomic coverage significantly hindered robust inference of melanocyte-to-keratinocyte interactions – an essential aspect when comparing the effects of co-cultured cell types between FibHSE and PmtHEE. To address this issue, we employed exFINDER^33^, which bypassed the need for a complete melanocyte cluster, enabling meaningful comparative analysis. Within the shared L1CAM-EZR-driven exSigNet, JUN was identified as the key transcription factor, serving as a rapid-response regulator of skin homeostasis that modulates the balance between keratinocyte proliferation and differentiation^53^, thereby contributing to barrier formation. Unlike broader and slower-acting signaling such as WNT, which is foundational for tissue patterning and long-term maintenance, JUN appears to act more acutely. Interestingly, JUN is not specific to PmtHEE; it is also broadly expressed in FibHSE keratinocytes (**Figure S7A**), raising the possibility that alternative ligand-receptor pairs may activate similar downstream targets to promote keratinocyte differentiation.

Indeed, ligands derived from fibroblasts or from the culture environment were found to activate JUN and other L1CAM-EZR-associated transcription factors, exerting similar regulatory effects on keratinocytes (**Figure S7C-D**). Nonetheless, despite shared TF executors in PmtHEE and FibHSE, functional enrichment analysis revealed model-specific transcriptional features (**Figure S7B**). Keratinocytes in FibHSE exhibited a more basal-like signature, with enrichment in ribosome biogenesis pathways, while those in PmtHEE demonstrated a suprabasal-like profile enriched in sphingolipid metabolic processes – functions closely linked to skin barrier formation and maintenance ^54^. These results suggest that co-cultured cell types distinctly influence epidermal differentiation trajectories, underscoring the importance of model context in skin bioengineering studies.

In light of this, numerous studies have demonstrated that keratinocyte-fibroblast crosstalk plays a critical role in regulating melanocyte distribution, survival, and melanin synthesis ^5,25,55,56^. Our melanocyte-containing PmtHEE model serves as a reductionist system to mimic skin pigmentation. However, the absence of a dermal compartment likely affects the experimental outcomes, as it omits the complex cellular communication between epidermal and dermal layers. This limitation may partly explain the observation of only a partial melanocyte-to-keratinocyte exSigNet in PmtHEE, along with the imbalance in cell proportions that reflects an enhanced differentiation profile. The development of fibroblast-integrated pigmented skin models may better recapitulate *in vivo*–like cell-cell interactions and downstream signaling pathways, potentially mitigating the disproportion between basal and suprabasal cell populations.

In summary, our study provides a comprehensive characterization of a widely used pigmented epidermal *in vitro* model. Comparative analyses revealed distinct differentiation capacities between fibroblast- and melanocyte-containing skin equivalents, both of which serve as valuable systems for *in vitro* skin modeling. Although these models form similarly stratified epidermis at the morphological level, their cellular behaviors diverge based on co-cultured cell types. These findings underscore the importance of single-cell resolution analysis and careful selection of *in vitro* skin models for their future applications.

## Methods

### Cell culture

Human primary epidermal melanocytes from a darkly pigmented donor (#C2025C, Thermo Fisher Scientific, Waltham, MA, USA) were cultured in melanocyte growth medium 2 (#C-24300, PromoCell, Heidelberg, Germany) until reaching 80% of confluence. Human primary epidermal keratinocytes (#C-12001; PromoCell, Heidelberg, Germany) were maintained in keratinocyte growth medium 2 (#C-20011, PromoCell, Heidelberg, Germany) up to 90% of confluence. Cells were kept at 37 °C in a 5% CO₂ humidified incubator.

### Pigmented human epidermal equivalent generation

For epidermal reconstruction, human primary epidermal melanocytes and keratinocytes were seeded at a density of 5×10⁵ cells/cm² onto 0.47 cm² inserts with a polycarbonate membrane (pore size: 0.4 µm; #140620, Thermo Fisher Scientific, Waltham, MA, USA) at a ratio of 1:20, respectively. Then, inserts were placed in 24-well plate filled up with 1.5 mL in the lower compartment and 500 μl in the upper compartment of coculture medium composed of 95% of EpiLife medium (#MEPI500CA, Thermo Fisher Scientific, Waltham, MA, USA) with additionally added 1.5 mM CaCl2 and human keratinocyte growth supplement (#S0015, Thermo Fisher Scientific, Waltham, MA, USA) and of 5% of melanocyte growth medium 2. After 72 h of incubation at 37 °C in a humidified atmosphere containing 5% CO₂, seeded cell cultures were raised to air-liquid interface (ALI) by removing the medium from the upper compartment. The medium in the lower compartment was changed to coculture medium additionally supplemented with 10 ng/ml keratinocyte growth factor (Sigma Aldrich, St. Louis, MO, USA), 50 µg/ml of vitamin C (Sigma Aldrich, St. Louis, MO, USA), and 50 µg/ml of gentamicin (Sigma Aldrich, St. Louis, MO, USA). Cell cultures were maintained at ALI for 14 days at 37°C in 5% CO₂ and humidified atmosphere. Medium was changed every 2-3 days.

### Histological analysis

The reconstructed tissues were fixed with 4% PFA for 24h at 4 °C. Following the fixation process, samples were dehydrated with gradually increasing concentrations of ethanol, concluding with xylene and embedding in paraffin. Subsequently, 5 µm-thick tissue sections were prepared and placed onto microscope glass slides (VWR, Mississauga, ON, USA). Later, samples underwent deparaffinization and rehydration with xylene and diminishing concentrations of ethanol, finishing with water. Hematoxylin and Eosin staining was performed following standard procedures. Samples were stained with Harris’s hematoxylin (VWR, Mississauga, ON, USA) for 3 minutes and Eosin Y (Sigma Aldrich, St. Louis, MO, USA) for 3 min. Fontana-Masson staining was performed using a commercially available kit (#ab150669; Abcam, Cambridge, UK), following the manufactureŕs instructions. Samples were mounted with DPX mounting medium (Sigma Aldrich, St. Louis, MO, USA) and visualized using a light microscope (Leica, Wetzlar, Germany) under 20x objective lens.

### Immunofluorescence staining

Immunolabeling was performed in the deparaffinized slides prepared for histological analysis described above. For keratin 14 and S100B detection, heat mediated antigen retrieval with sodium citrate buffer (pH 6.0) at 95 °C for 20 min was performed. Subsequently, samples were washed with 0.1 M glycine buffer. For blocking, the slides were immersed in 0.2% BSA in PBS for 1 h. Then, samples were incubated at 4°C overnight with the primary antibody anti-cytokeratin 14 (1:200, #ab7800, mouse, monoclonal, IgG3, Abcam) or anti-S100B (1:200, #ab52642, rabbit, monoclonal, IgG; Abcam). For keratin 10, involucrin and loricrin detection, tissue sections were rinsed with 0.1 M glycine buffer, followed by blocking and permeabilization with 0.2% BSA and 0.02% Triton X-100 in PBS. Afterwards, samples were incubated for 1 h with primary antibodies anti-cytokeratin 10 (1:200, #ab76318, rabbit, monoclonal, IgG; Abcam), anti-involucrin (1:200, #I9018, mouse, monoclonal, IgG1; Sigma Aldrich) or anti-loricrin (1:200, #ab85679, rabbit, polyclonal, IgG, abcam, Cambridge, UK). Samples were incubated for 45 min with the secondary antibody anti-rabbit Alexa Fluor 488 (1:200, #A21206, donkey, polyclonal, IgG; Thermo Fisher Scientific) or anti-mouse Alexa Fluor 594 (1:200, #A21203, donkey, polyclonal, IgG; Thermo Fisher Scientific). Slides were mounted with Fluoromount G® mounting medium with DAPI (SouthernBiotech, Birmingham, AL, USA). Fluorescent images were taken using a Nikon Eclipse 90i microscope at 20X magnification.

### Transmission Electron Microscopy (TEM)

Samples were fixed in 2% paraformaldehyde with 2.5% glutaraldehyde in cacodylate 0.1 M overnight at 4°C. Then, tissues were post-fixed with 1% osmium tetroxide with 0.8% potassium ferrocyanide for 2 h and subsequently dehydrated with a series of increasing acetone concentrations. After embedding in epon resin (EMS, Hatfield, PA, USA), samples polymerized at 70°C for 48 h. Sections of 70 nm in thickness were obtained with a Reichert-Jung Ultracut E ultramicrotome. Next, samples were double stained with 1% uranyl acetate for 45 minutes (Sigma Aldrich, Steinheim, Germany) and 3% lead citrate (Leica Biosystems, Nussloch, Germany) for 5 min at 20°C in an automatic contrasting system Leica EM AC20. Samples were viewed by a TEM Hitachi H-7000 transmission electron microscope (JEOL, Akishima, Tokyo, Japan), and imaged at a voltage of 75 kV.

### Tissue dissociation and single-cell isolation

On the last day of culture, 15 PmtHEE samples were placed in one Eppendorf tube with sCelLive™ Tissue Preservation Solution (cat. nr. 10100611, Singleron Biotechnologies GmbH, Cologne, Germany), as previously described^57^ and stored at 4°C until further processing. Later, tissues were washed with 1x Hank’s Balanced Saline Solution (HBSS, Thermo Fisher Scientific) and dissociated with ophthalmic scissors to finely cut pieces of 1-2 mm. The pieces were digested in 4.5 mL Skin Tissue Dissociation Solution B (cat. nr. 1200050037, Singleron Biotechnologies) at 37°C for 2 x 25 min with continuous agitation on a thermal shaker at 350 rpm. The state of dissociation was checked at regular intervals under a light microscope. Following the first step of digestion, the suspension was filtered using a 100-µm sterile strainer (cat. nr. 130-098-463, Miltenyi, Bergisch Gladbach, Germany) adding 0.04% PBS-BSA. The remaining undissociated tissue was digested in 4mL Skin Tissue Dissociation Solution C (cat. nr. 1200050031, Singleron Biotechnologies) at 37°C for 2 x 25 min with continuous agitation on a thermal shaker at 350 rpm. The suspension was filtered again using a 100-µm sterile strainer. The cells were centrifuged at 350 x g for 5 min at 4°C and the cell pellets were pooled and resuspended in 250mL of PBS. The cells were counted using acridine orange/propidium iodide with a Luna FX7 automated cell counter (Logos Biosystems, Villeneuve d’Ascq, France).

### Single cell library preparation and sequencing

The resulting single-cell suspension was directly loaded onto a microfluidic SCOPE-chip and processed for scRNA-seq library preparation according to the manufacturer’s protocol (GEXSCOPE^TM^ Single Cell RNA-seq Kit, Singleron Biotechnologies). The ultimately constructed and purified library was sequenced on the Illumina NovaSeq X with a paired-end 150-bp approach at a depth of 1 GB per well. The CeleScoot pipeline (v1.0.0) was used for scRNA-seq data alignment and quantification, against the *Homo sapiens* GRCh38 (Ensembl release 99, https://ftp.ensembl.org/pub/release-99/fasta/homo_sapiens/).

### Single cell data processing

In addition to the data generated in this study, the following publicly available external single-cell datasets were obtained from the Gene Expression Omnibus (GEO) (https://www.ncbi.nlm.nih.gov/geo/), including GSE190695 (HSEs) ^19^ and GSE147482 (human neonatal foreskin) ^16^.

A total of 8 scRNA-seq datasets representing *in vivo* control (5 FsEpi samples) and multiple *in vitro* skin cultures (two FibHSE samples, and one PmtHEE sample) were processed using the Seurat single-cell analysis pipeline (v4.3.0)^58^ in R (v4.4.1). For the quality control, genes detected in <3 cells were removed first, and low-quality cells were further filtered based on sample-specific QC metrics:

- FsEpi samples: Cells with >500 and <5,000 or 6000 detected genes per cell, <30,000 or 50,000 UMI counts per cell, and <15% or 20% mitochondrial gene expression were retained;
- FibHSE samples: Cells with >600 and <7,000 detected genes per cell, <70,000 UMI counts per cell, and <20% mitochondrial gene expression were retained;
- PmtHEE sample: Cells with >200 and <5,000 detected genes per cell, <40,000 UMI counts per cell, and <20% mitochondrial gene expression were included in the analysis. After filtering, data in each cell were normalized, the 2,000 most variable genes were identified, and the expression levels of these genes were scaled before performing PCA in variable gene space. Next, 30 principal components were used for graph-based clustering and UMAP dimensionality reduction was computed. All steps were performed using functions implemented in the Seurat package (*NormalizeData, FindVariableFeatures, ScaleData, RunPCA, FindNeighbours, FindClusters, RunUMAP*) with default parameters, except where mentioned.

To identify potential doublets, we employed DoubletFinder(v2.0)^59^ and scDblFinder(v1.18.0)^60^. Doublets predicted by both algorithms and annotated as *’doublet:Doublet’* were selectively removed to ensure accurate results while preserving as many cells as possible. Only a small number of doublets were detected in the dataset, and no doublet-specific clusters were predicted by either algorithm.

The processed and combined list objects were subsequently used for batch effect correction and dataset integration using Seurat’s canonical correlation analysis (CCA) integration method^61^, with parameters set to *nfeatures = 2,000, anchor.features = 2,000,* and *dims = 1:30*. To determine the optimal clustering resolution, we evaluated multiple resolutions (0.5, 0.7, and 0.9) based on Seurat’s built-in cell cycle scoring, ROGUE statistic^62^, and the biologically meaningful distribution of cells across clusters inferred from canonical marker gene expression. A resolution of 0.7 was selected for the final UMAP visualization. Cell type identities were assigned to each cluster based on the expression of established skin lineage marker genes.

### Identification of differentially expressed genes

Differentially expressed genes (DEGs) between keratinocytes from FibHSE and PmtHEE were identified using the *FindMarkers()* function in Seurat, with the following parameters: *min.pct = 0.25, logfc.threshold = 0.25* and *pseudocount.use = 1*. The non-parametric Wilcoxon rank-sum test was applied to compute p-values, and Bonferroni correction was used to adjust p-values across all genes in the dataset. The top 200 DEGs for each condition were selected based on an adjusted p-value threshold of < 0.01 and ranked by average log2 fold change (from highest to lowest). These DEGs were subsequently used as input for functional enrichment analysis.

### Biological process functional enrichment analysis

Gene Ontology (GO) enrichment analysis was performed using the *clusterProfiler* package^63^. The *bitr()* function was first used to convert gene symbols to Entrez IDs, referencing the *org.Hs.eg.db* database (version 3.18.0)^64^. GO enrichment was then conducted using the *enrichGO()* function with the *parameters ont = "BP", pAdjustMethod = "BH",* and *qvalueCutoff = 0.05*. The top 5 or 10 most significantly enriched biological process categories were selected for downstream visualization.

### Cell lineage and pseudotime inference

We inferred cell lineage trajectories within the keratinocyte compartment to capture the transition from basal to terminal differentiation states using *Slingshot* (v2.7.0)^31^, with default parameters. A minimum spanning tree (MST) was constructed between clusters based on their centroids in the reduced-dimensional embedding space. For each inferred lineage, pseudotime values were calculated as the arc-length along a fitted principal curve, originating from the root cluster, which was automatically determined by the algorithm.

### Cell-cell communication analysis

Global differences in cell-cell communication between PmtHEE and FsEpi were evaluated using *CellChat* (v2.1.2)^32^. Specific interactions, including melanocyte-to-keratinocyte and basal keratinocyte-to-melanocyte signaling, as well as external signaling networks (exSigNets) targeting recipient cells, were inferred using *exFINDER* (v1.0.0)^33^. For downstream analysis, the top 10 marker genes of BAS, SPN, and GRN keratinocyte subtypes were selected as keratinocyte target genes, while the top 10 marker genes of melanocytes were used as their corresponding target gene set. The *percentile = c(0.5, 0.75, 0.90)* argument was applied to define the target gene expression thresholds, corresponding to the median, upper quartile, and 90th percentile of the gene expression distribution, respectively. Expression thresholds for low and high expression in target keratinocyte cells were defined as 0.03 and 0.45 in FsEpi, and 0.05 and 0.6 in FibHSE. For melanocyte target cells in FsEpi, low and high expression thresholds were set at 0.03 and 0.5, respectively, while thresholds for keratinocyte targets in PmtHEE were set at 0.02 and 0.3.

## Acknowledgments

S.X.A. and Y.Y.J acknowledge the support of the National Science Foundation (CBET2134916). J.B. was supported by a Doctorats Industrials grant from the Generalitat de Catalunya (2020 DI 00082).

## Author contributions

Conceptualization and methodology, Y. Y. J., J. B., M. P.-A., and S. X. A.; investigation, Y. Y. J., J. B., M. P.-A., and S. X. A.; formal analysis, Y. Y. J., J. B., M. P.-A., and S. X. A.; writing – original draft, review & editing, Y. Y. J., J. B., M. P.-A., and S. X. A; funding acquisition and supervision, M. P.-A., and S. X. A. All authors discussed the results and commented on the manuscript.

---

**Figure S1.**
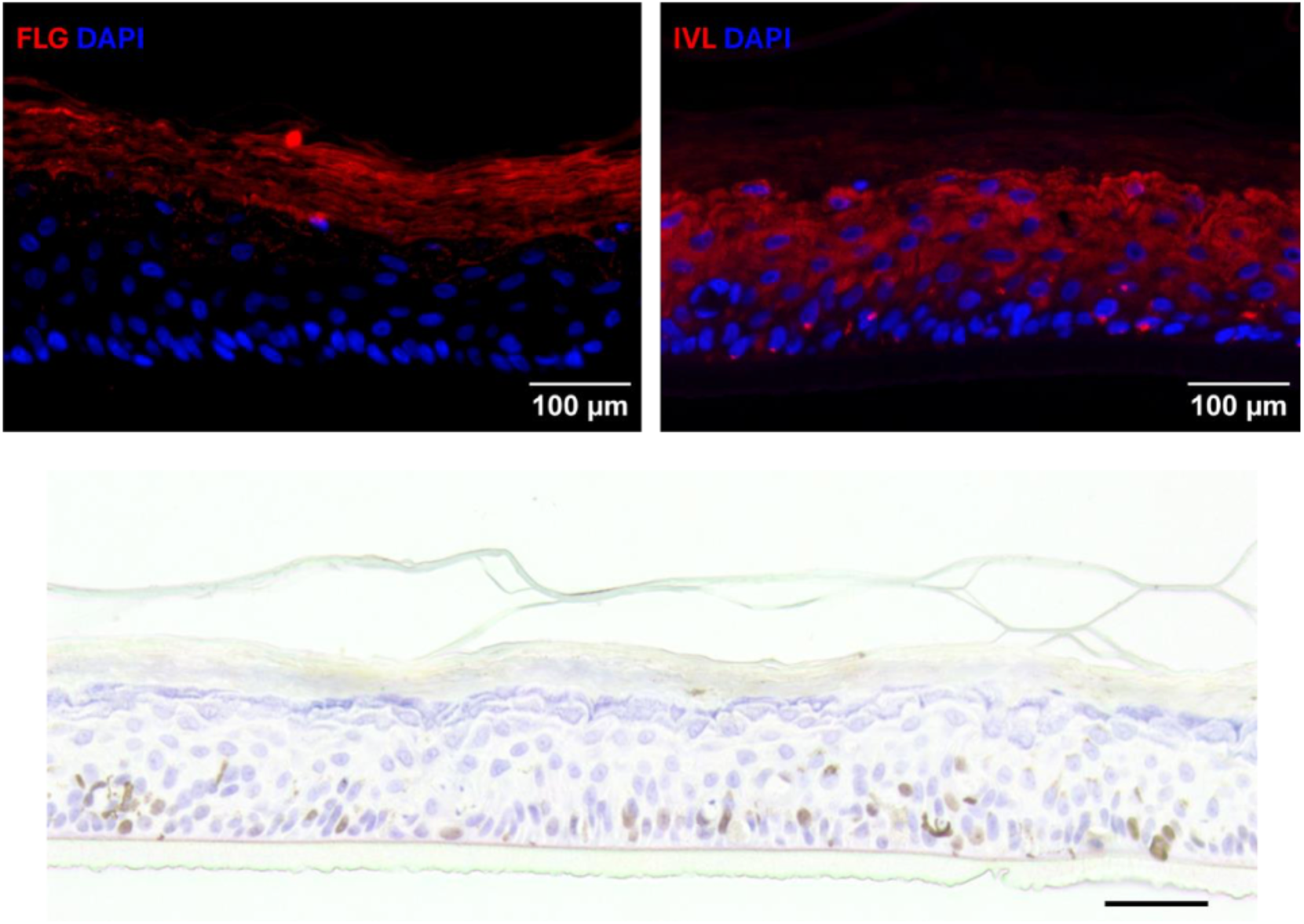
Representative images showing the expression of marker proteins detected by immunofluorescence (IF, upper panels) and immunohistochemistry (IHC, lower panel). FLG (upper left), IVL (upper right), and MKI67 (bottom) were analyzed to visualize their spatial distribution within the tissue. Cell nuclei were counterstained with DAPI. Scale bars: 100 µm (IF) and 50 µm (IHC).

**Figure S2.**
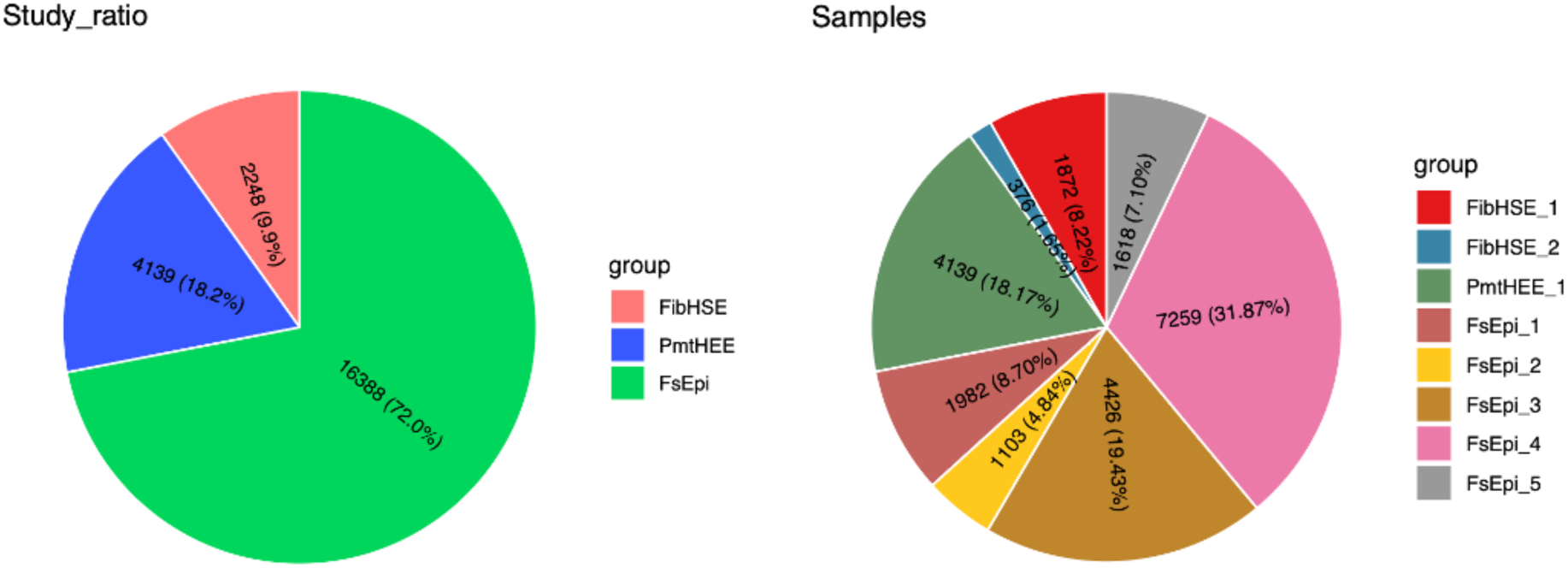
Pie chart illustrating the composition of scRNA-seq datasets.

**Figure S3.**
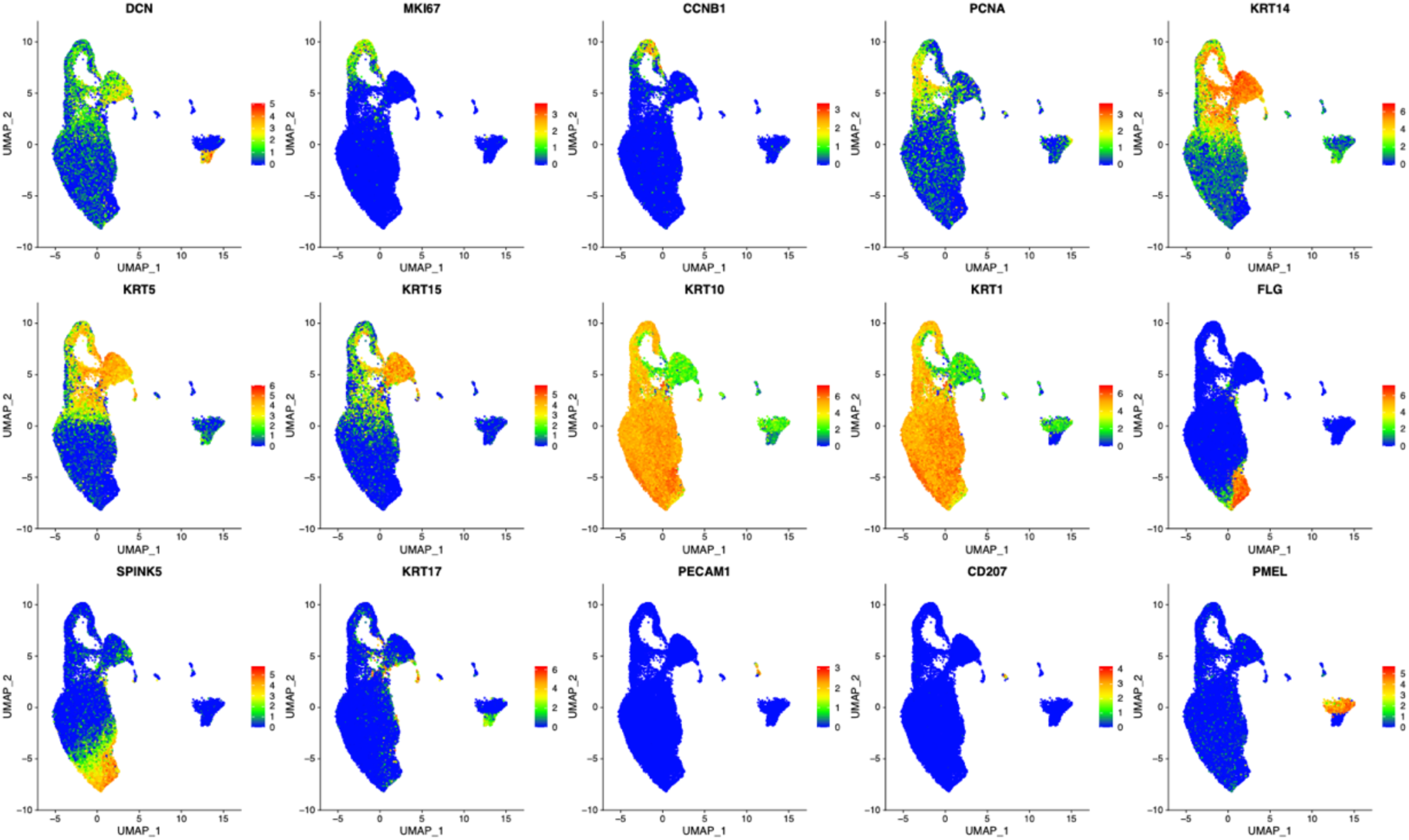
Feature plots showing the expression of canonical skin marker genes across cell populations within the integrated Seurat object.

**Figure S4.**
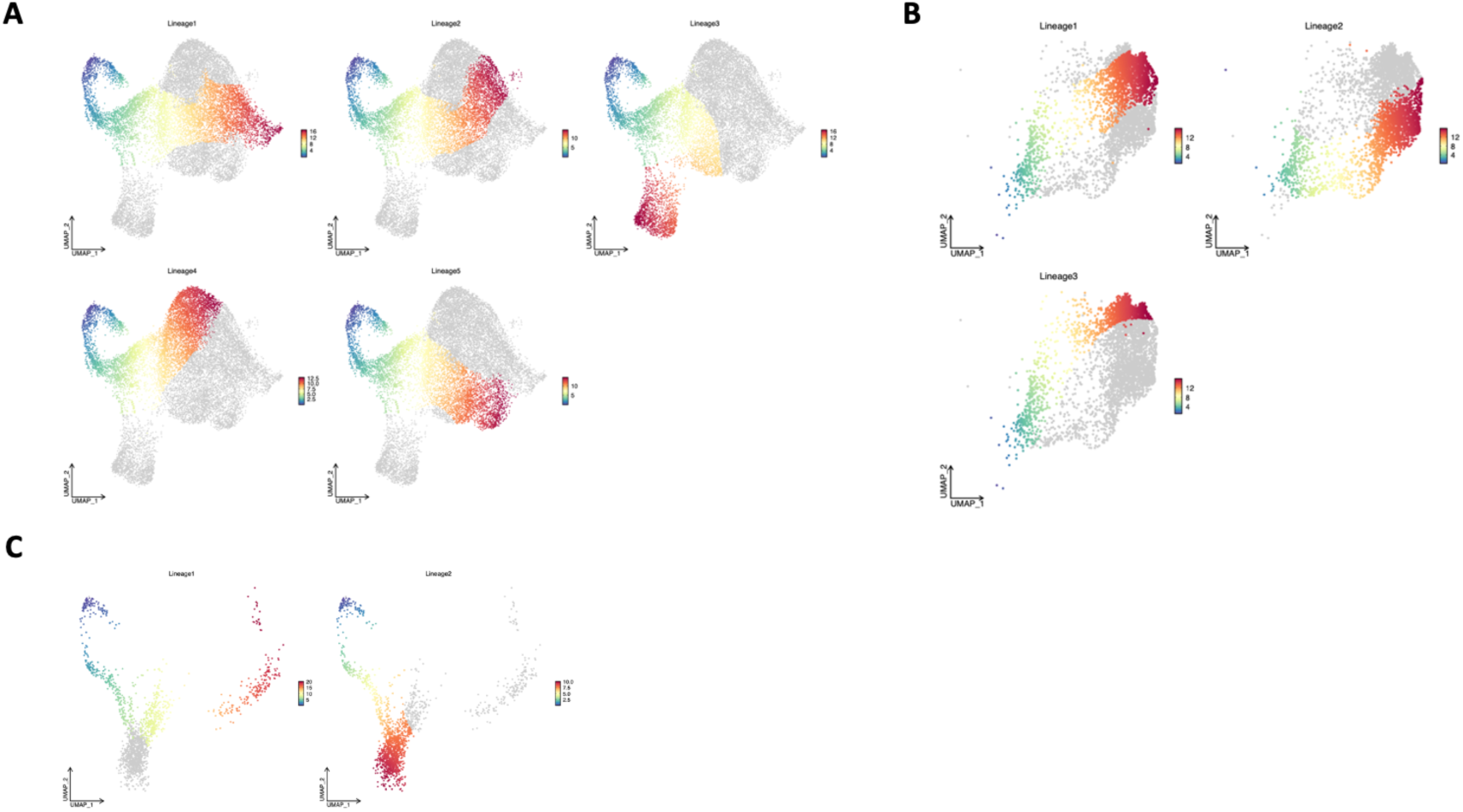
Pseudotime inference plots showing stable estimates of the underlying cell-level pseudotime variable for each lineage. Panels display results for (A) FsEpi, (B) PmtHEE, and (C) FibHSE models.

**Figure S5.**
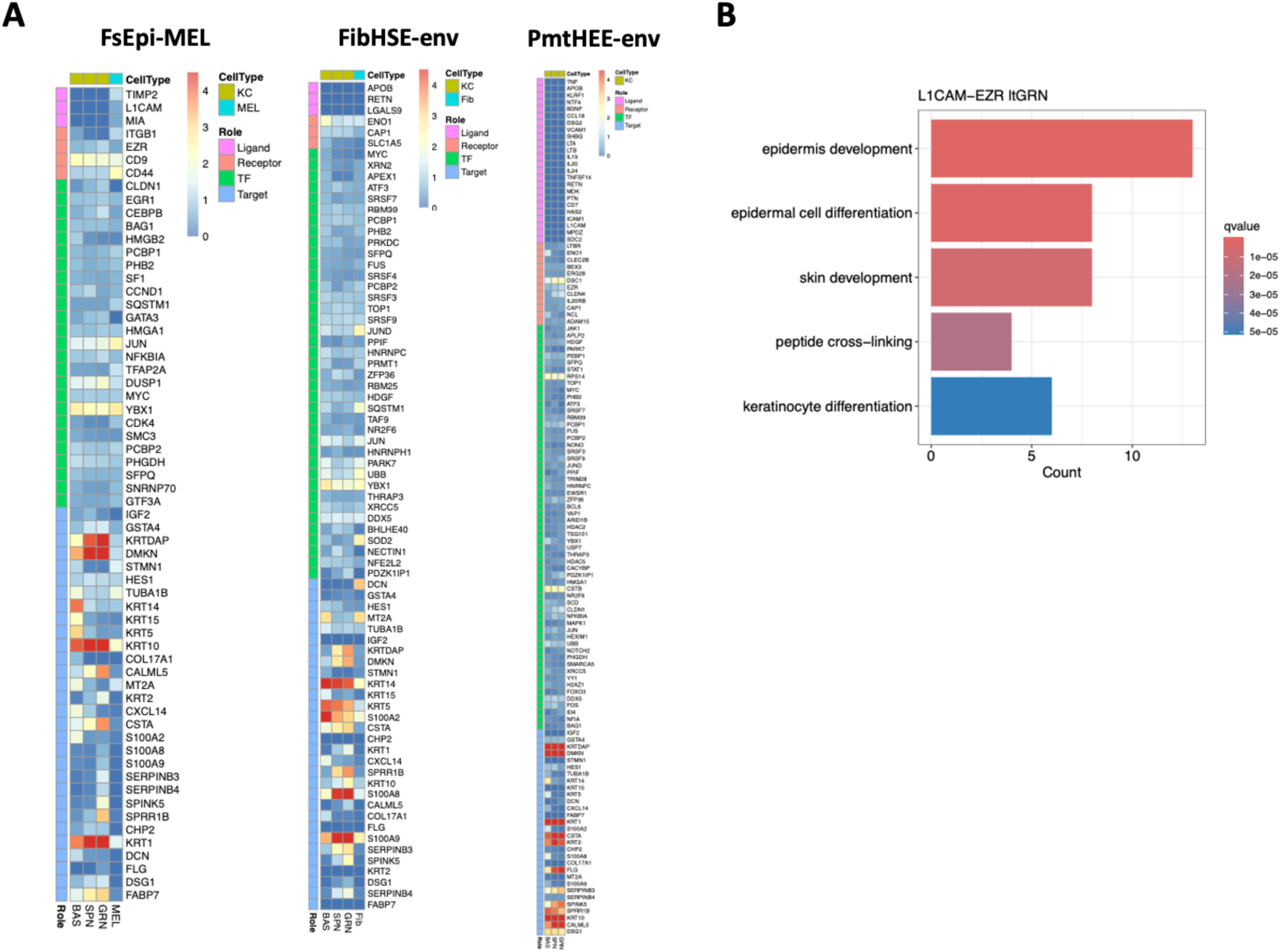
(A) Heatmap showing the expression levels of components from predicted exSigNets derived from selected sources under FsEpi, FibHSE, and PmtHEE conditions. (B) Functional enrichment analysis of biological processes for the shared L1CAM-EZR-driven exSigNet.

**Figure S6.**
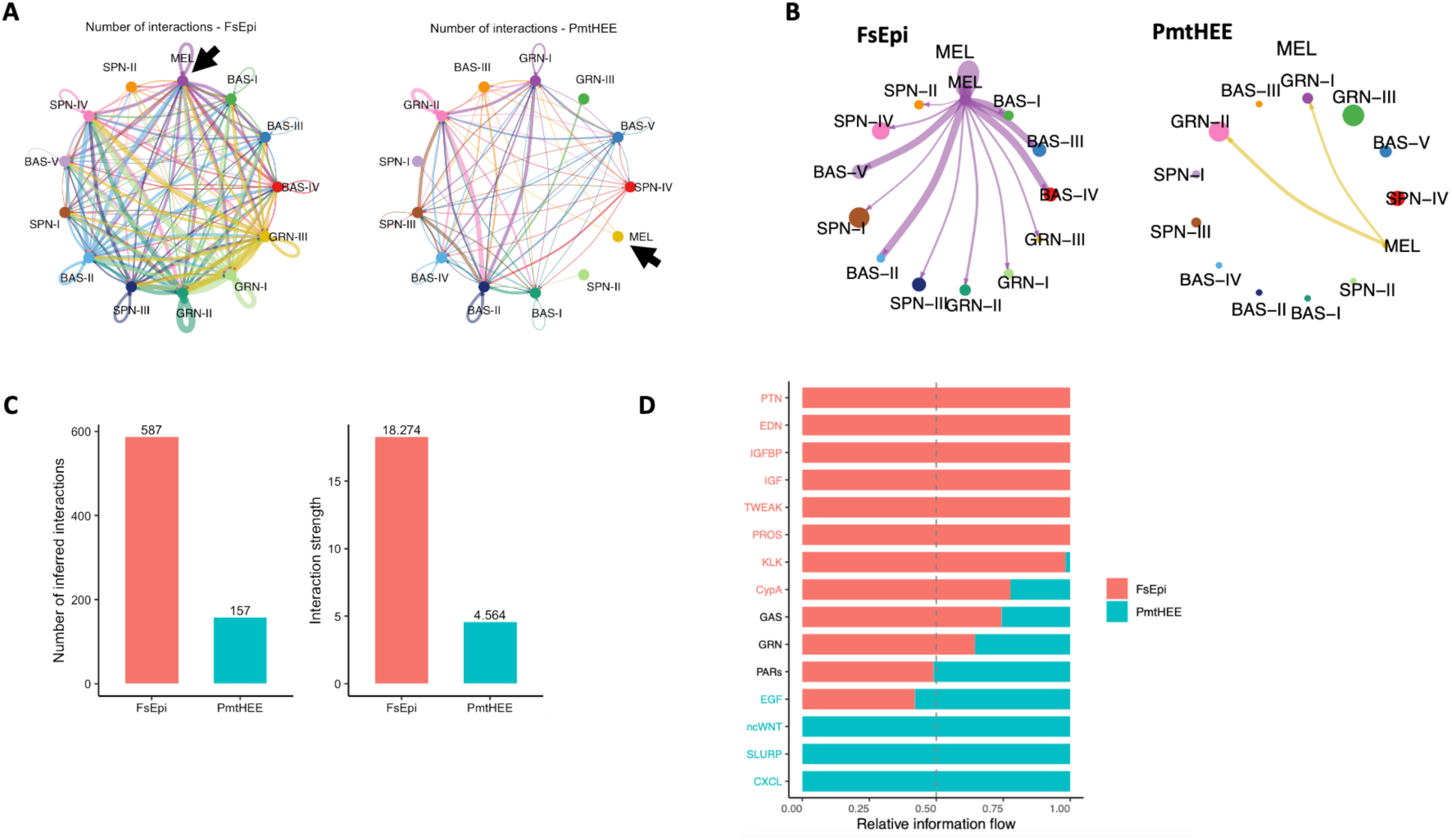
Comparative CellChat analysis of intercellular communication in FsEpi and PmtHEE skin models. (A) Network plot showing the total number of significant ligand-receptor interactions between cell populations in each model. (B) The outgoing signaling strength of melanocyte-derived communication across cell types. (C) Bar plot comparing the total number of cell-cell interactions detected in FsEpi and PmtHEE. (D) Bar plot of relative information flow for key signaling pathways, comparing their contributions in FsEpi (coral red) versus PmtHEE (teal blue).

**Figure S7.**
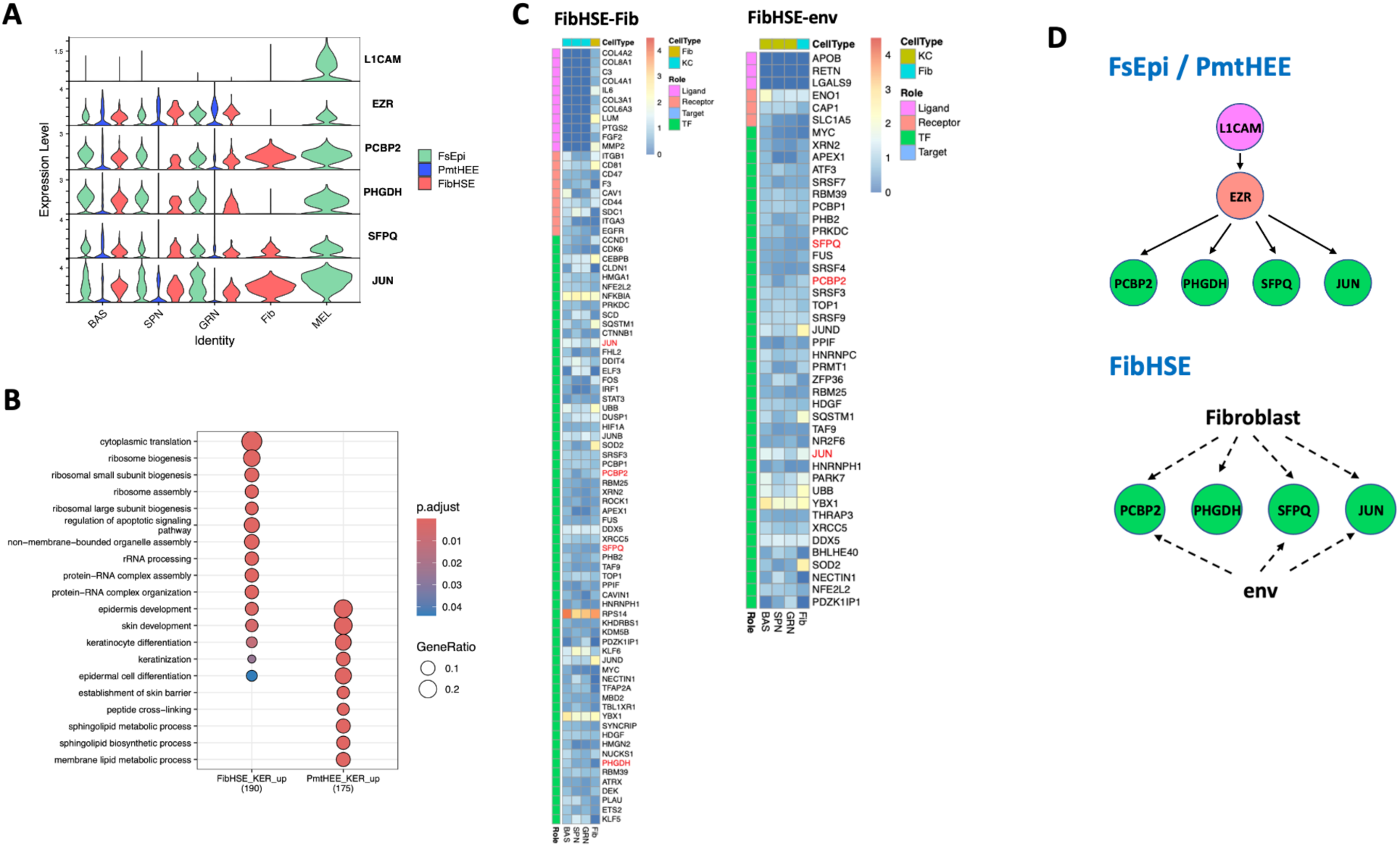
Alternative exSigNets targeting keratinocytes identified in the FibHSE model. (A) Violin plot showing the expression of ligand-receptor-transcription factor components from the shared L1CAM-EZR-driven exSigNet, predicted between FsEpi and PmtHEE. The FibHSE model is now included to enable comprehensive comparison across all conditions. (B) Functional enrichment analysis of biological processes in keratinocytes from FibHSE and PmtHEE. (C) Heatmap showing the expression levels of components within the fibroblast- and environment-specific exSigNets identified in the FibHSE model. Due to substantial overlap of target genes with those in Figure 6C, only ligand-receptor-transcription factor components are presented here. Transcription factors shared with the FsEpi/PmtHEE-derived L1CAM-EZR-driven exSigNet are highlighted in red. ’FibHSE_Fib’ represents ligands derived from fibroblasts in the FibHSE model; ’FibHSE_env’ refers to ligands from the external environment in the FibHSE model. Keratinocytes were used as the target cells in all analyses. (D) Schematic illustrating the origin of alternative exSigNets in the FibHSE model that bypass the L1CAM-EZR-driven exSigNet, in contrast to the melanocyte-to-keratinocyte signaling observed in FsEpi and PmtHEE.

## References

1. Abdel-Naser, M.B., Nikolakis, G., and Zouboulis, C.C. (2023). Preservation of epidermal melanocyte integrity in an ex vivo co-culture skin model with sebocytes. Exp. Dermatol. 32, 1063–1071. 10.1111/exd.14813.

2. Casalou, C., Moreiras, H., Mayatra, J.M., Fabre, A., and Tobin, D.J. (2022). Loss of ‘Epidermal Melanin Unit’ Integrity in Human Skin During Melanoma-Genesis. Front. Oncol. 12, 878336. 10.3389/fonc.2022.878336.

3. Kaidbey, K.H., Agin, P.P., Sayre, R.M., and Kligman, A.M. (1979). Photoprotection by melanin—a comparison of black and Caucasian skin. J. Am. Acad. Dermatol. 1, 249–260. 10.1016/S0190-9622(79)70018-1.

4. Chen, Y.-Y., Liu, L.-P., Zhou, H., Zheng, Y.-W., and Li, Y.-M. (2022). Recognition of Melanocytes in Immuno-Neuroendocrinology and Circadian Rhythms: Beyond the Conventional Melanin Synthesis. Cells 11, 2082. 10.3390/cells11132082.

5. Bastonini, E., Kovacs, D., and Picardo, M. (2016). Skin Pigmentation and Pigmentary Disorders: Focus on Epidermal/Dermal Cross-Talk. Ann. Dermatol. 28, 279–289. 10.5021/ad.2016.28.3.279.

6. Slominski, R.M., Sarna, T., Płonka, P.M., Raman, C., Brożyna, A.A., and Slominski, A.T. (2022). Melanoma, Melanin, and Melanogenesis: The Yin and Yang Relationship. Front. Oncol. 12, 842496. 10.3389/fonc.2022.842496.

7. Reijnders, C.M.A., van Lier, A., Roffel, S., Kramer, D., Scheper, R.J., and Gibbs, S. (2015). Development of a Full-Thickness Human Skin Equivalent In Vitro Model Derived from TERT-Immortalized Keratinocytes and Fibroblasts. Tissue Eng. Part A 21, 2448–2459. 10.1089/ten.tea.2015.0139.

8. Topol, B.M., Haimes, H.B., Dubertret, L., and Bell, E. (1986). Transfer of melanosomes in a skin equivalent model in vitro. J. Invest. Dermatol. 87, 642–647. 10.1111/1523-1747.ep12456314.

9. Hofmann, E., Schwarz, A., Fink, J., Kamolz, L.-P., and Kotzbeck, P. (2023). Modelling the Complexity of Human Skin In Vitro. Biomedicines 11, 794. 10.3390/biomedicines11030794.

10. Hosseini, M., Koehler, K.R., and Shafiee, A. (2022). Biofabrication of Human Skin with Its Appendages. Adv. Healthc. Mater. 11, 2201626. 10.1002/adhm.202201626.

11. Jia, Y.Y., and Atwood, S.X. (2024). Diversity of human skin three-dimensional organotypic cultures. Curr. Opin. Genet. Dev. 89, 102275. 10.1016/j.gde.2024.102275.

12. Wang, X.-Y., Jia, Q.-N., Li, J., and Zheng, H.-Y. (2024). Organoids as Tools for Investigating Skin Aging: Mechanisms, Applications, and Insights. Biomolecules 14, 1436. 10.3390/biom14111436.

13. Zamudio Díaz, D.F., Busch, L., Kröger, M., Klein, A.L., Lohan, S.B., Mewes, K.R., Vierkotten, L., Witzel, C., Rohn, S., and Meinke, M.C. (2024). Significance of melanin distribution in the epidermis for the protective effect against UV light. Sci. Rep. 14, 3488. 10.1038/s41598-024-53941-0.

14. Glover, J.D., Sudderick, Z.R., Shih, B.B.-J., Batho-Samblas, C., Charlton, L., Krause, A.L., Anderson, C., Riddell, J., Balic, A., Li, J., et al. (2023). The developmental basis of fingerprint pattern formation and variation. Cell 186, 940–956.e20. 10.1016/j.cell.2023.01.015.

15. Ober-Reynolds, B., Wang, C., Ko, J.M., Rios, E.J., Aasi, S.Z., Davis, M.M., Oro, A.E., and Greenleaf, W.J. (2023). Integrated single-cell chromatin and transcriptomic analyses of human scalp identify gene-regulatory programs and critical cell types for hair and skin diseases. Nat. Genet. 55, 1288–1300. 10.1038/s41588-023-01445-4.

16. Wang, S., Drummond, M.L., Guerrero-Juarez, C.F., Tarapore, E., MacLean, A.L., Stabell, A.R., Wu, S.C., Gutierrez, G., That, B.T., Benavente, C.A., et al. (2020). Single cell transcriptomics of human epidermis identifies basal stem cell transition states. Nat. Commun. 11, 4239. 10.1038/s41467-020-18075-7.

17. Gopee, N.H., Winheim, E., Olabi, B., Admane, C., Foster, A.R., Huang, N., Botting, R.A., Torabi, F., Sumanaweera, D., Le, A.P., et al. (2024). A prenatal skin atlas reveals immune regulation of human skin morphogenesis. Nature, 1–11. 10.1038/s41586-024-08002-x.

18. Lee, J., Rabbani, C.C., Gao, H., Steinhart, M.R., Woodruff, B.M., Pflum, Z.E., Kim, A., Heller, S., Liu, Y., Shipchandler, T.Z., et al. (2020). Hair-bearing human skin generated entirely from pluripotent stem cells. Nature 582, 399–404. 10.1038/s41586-020-2352-3.

19. Stabell, A.R., Lee, G.E., Jia, Y., Wong, K.N., Wang, S., Ling, J., Nguyen, S.D., Sen, G.L., Nie, Q., and Atwood, S.X. (2023). Single-cell transcriptomics of human-skin-equivalent organoids. Cell Rep. 42, 112511. 10.1016/j.celrep.2023.112511.

20. Hatzfeld, M., Keil, R., and Magin, T.M. (2017). Desmosomes and Intermediate Filaments: Their Consequences for Tissue Mechanics. Cold Spring Harb. Perspect. Biol. 9, a029157. 10.1101/cshperspect.a029157.

21. Bento-Lopes, L., Cabaço, L.C., Charneca, J., Neto, M.V., Seabra, M.C., and Barral, D.C. (2023). Melanin’s Journey from Melanocytes to Keratinocytes: Uncovering the Molecular Mechanisms of Melanin Transfer and Processing. Int. J. Mol. Sci. 24, 11289. 10.3390/ijms241411289.

22. Hall, M.J., Lopes-Ventura, S., Neto, M.V., Charneca, J., Zoio, P., Seabra, M.C., Oliva, A., and Barral, D.C. (2022). Reconstructed human pigmented skin/epidermis models achieve epidermal pigmentation through melanocore transfer. Pigment Cell Melanoma Res. 35, 425–435. 10.1111/pcmr.13039.

23. Bowman, S.L., and Marks, M.S. (2018). Shining a Light on Black Holes in Keratinocytes. J. Invest. Dermatol. 138, 486–489. 10.1016/j.jid.2017.11.002.

24. Arnette, C.R., Roth-Carter, Q.R., Koetsier, J.L., Broussard, J.A., Burks, H.E., Cheng, K., Amadi, C., Gerami, P., Johnson, J.L., and Green, K.J. (2020). Keratinocyte cadherin desmoglein 1 controls melanocyte behavior through paracrine signaling. Pigment Cell Melanoma Res. 33, 305–317. 10.1111/pcmr.12826.

25. Goncalves, K., De Los Santos Gomez, P., Costello, L., Smith, L., Mead, H., Simpson, A., and Przyborski, S. (2022). Investigation into the effect of skin tone modulators and exogenous stress on skin pigmentation utilizing a novel bioengineered skin equivalent. Bioeng. Transl. Med. 8, e10415. 10.1002/btm2.10415.

26. Roger, M., Fullard, N., Costello, L., Bradbury, S., Markiewicz, E., O’Reilly, S., Darling, N., Ritchie, P., Määttä, A., Karakesisoglou, I., et al. (2019). Bioengineering the microanatomy of human skin. J. Anat. 234, 438–455. 10.1111/joa.12942.

27. Cheng, J.B., Sedgewick, A.J., Finnegan, A.I., Harirchian, P., Lee, J., Kwon, S., Fassett, M.S., Golovato, J., Gray, M., Ghadially, R., et al. (2018). Transcriptional Programming of Normal and Inflamed Human Epidermis at Single-Cell Resolution. Cell Rep. 25, 871–883. 10.1016/j.celrep.2018.09.006.

28. Xu, Z., Chen, D., Hu, Y., Jiang, K., Huang, H., Du, Y., Wu, W., Wang, J., Sui, J., Wang, W., et al. (2022). Anatomically distinct fibroblast subsets determine skin autoimmune patterns. Nature 601, 118–124. 10.1038/s41586-021-04221-8.

29. Zou, Z., Long, X., Zhao, Q., Zheng, Y., Song, M., Ma, S., Jing, Y., Wang, S., He, Y., Esteban, C.R., et al. (2021). A Single-Cell Transcriptomic Atlas of Human Skin Aging. Dev. Cell 56, 383–397.e8. 10.1016/j.devcel.2020.11.002.

30. Crow, M., Paul, A., Ballouz, S., Huang, Z.J., and Gillis, J. (2018). Characterizing the replicability of cell types defined by single cell RNA-sequencing data using MetaNeighbor. Nat. Commun. 9, 884. 10.1038/s41467-018-03282-0.

31. Street, K., Risso, D., Fletcher, R.B., Das, D., Ngai, J., Yosef, N., Purdom, E., and Dudoit, S. (2018). Slingshot: cell lineage and pseudotime inference for single-cell transcriptomics. BMC Genomics 19, 477. 10.1186/s12864-018-4772-0.

32. Jin, S., Guerrero-Juarez, C.F., Zhang, L., Chang, I., Ramos, R., Kuan, C.-H., Myung, P., Plikus, M.V., and Nie, Q. (2021). Inference and analysis of cell-cell communication using CellChat. Nat. Commun. 12, 1088. 10.1038/s41467-021-21246-9.

33. He, C., Zhou, P., and Nie, Q. (2023). exFINDER: identify external communication signals using single-cell transcriptomics data. Nucleic Acids Res. 51, e58. 10.1093/nar/gkad262.

34. van der Maten, M., Reijnen, C., Pijnenborg, J.M.A., and Zegers, M.M. (2019). L1 Cell Adhesion Molecule in Cancer, a Systematic Review on Domain-Specific Functions. Int. J. Mol. Sci. 20, 4180. 10.3390/ijms20174180.

35. Bell, S., Degitz, K., Quirling, M., Jilg, N., Page, S., and Brand, K. (2003). Involvement of NF-kappaB signalling in skin physiology and disease. Cell. Signal. 15, 1–7. 10.1016/s0898-6568(02)00080-3.

36. Eckert, R.L., Adhikary, G., Young, C.A., Jans, R., Crish, J.F., Xu, W., and Rorke, E.A. (2013). AP1 Transcription Factors in Epidermal Differentiation and Skin Cancer. J. Skin Cancer 2013, 537028. 10.1155/2013/537028.

37. Bi, O., Anene, C.A., Nsengimana, J., Shelton, M., Roberts, W., Newton-Bishop, J., and Boyne, J.R. (2021). SFPQ promotes an oncogenic transcriptomic state in melanoma. Oncogene 40, 5192–5203. 10.1038/s41388-021-01912-4.

38. Mattaini, K.R., Sullivan, M.R., Lau, A.N., Fiske, B.P., Bronson, R.T., and Vander Heiden, M.G. (2019). Increased PHGDH expression promotes aberrant melanin accumulation. BMC Cancer 19, 723. 10.1186/s12885-019-5933-5.

39. Ren, C., Zhang, J., Yan, W., Zhang, Y., and Chen, X. (2016). RNA-binding Protein PCBP2 Regulates p73 Expression and p73-dependent Antioxidant Defense. J. Biol. Chem. 291, 9629–9637. 10.1074/jbc.M115.712125.

40. Bell, E., Ehrlich, H.P., Buttle, D.J., and Nakatsuji, T. (1981). Living tissue formed in vitro and accepted as skin-equivalent tissue of full thickness. Science 211, 1052–1054. 10.1126/science.7008197.

41. Pruniéras, M., Régnier, M., and Woodley, D. (1983). Methods for cultivation of keratinocytes with an air-liquid interface. J. Invest. Dermatol. 81, 28s–33s. 10.1111/1523-1747.ep12540324.

42. Cousin, I., Misery, L., de Vries, P., and Lebonvallet, N. (2023). Emergence of New Concepts in Skin Physiopathology through the Use of in vitro Human Skin Explants Models. Dermatology 239, 849–859. 10.1159/000533261.

43. de Groot, S.C., Ulrich, M.M.W., Gho, C.G., and Huisman, M.A. (2021). Back to the Future: From Appendage Development Toward Future Human Hair Follicle Neogenesis. Front. Cell Dev. Biol. 9, 661787. 10.3389/fcell.2021.661787.

44. Kim, J., Koo, B.-K., and Knoblich, J.A. (2020). Human organoids: model systems for human biology and medicine. Nat. Rev. Mol. Cell Biol. 21, 571–584. 10.1038/s41580-020-0259-3.

45. Lee, J., van der Valk, W.H., Serdy, S.A., Deakin, C., Kim, J., Le, A.P., and Koehler, K.R. (2022). Generation and characterization of hair-bearing skin organoids from human pluripotent stem cells. Nat. Protoc. 17, 1266–1305. 10.1038/s41596-022-00681-y.

46. Arredondo, J., Chernyavsky, A.I., Webber, R.J., and Grando, S.A. (2005). Biological effects of SLURP-1 on human keratinocytes. J. Invest. Dermatol. 125, 1236–1241. 10.1111/j.0022-202X.2005.23973.x.

47. Korbecki, J., Maruszewska, A., Bosiacki, M., Chlubek, D., and Baranowska-Bosiacka, I. (2022). The Potential Importance of CXCL1 in the Physiological State and in Noncancer Diseases of the Cardiovascular System, Respiratory System and Skin. Int. J. Mol. Sci. 24, 205. 10.3390/ijms24010205.

48. Lim, X., and Nusse, R. (2013). Wnt Signaling in Skin Development, Homeostasis, and Disease. Cold Spring Harb. Perspect. Biol. 5, a008029. 10.1101/cshperspect.a008029.

49. Florin, L., Maas-Szabowski, N., Werner, S., Szabowski, A., and Angel, P. (2005). Increased keratinocyte proliferation by JUN-dependent expression of PTN and SDF-1 in fibroblasts. J. Cell Sci. 118, 1981–1989. 10.1242/jcs.02303.

50. Nauroy, P., and Nyström, A. (2020). Kallikreins: Essential epidermal messengers for regulation of the skin microenvironment during homeostasis, repair and disease. Matrix Biol. Plus 6*–*7, 100019. 10.1016/j.mbplus.2019.100019.

51. Liu, Q., Xiao, S., and Xia, Y. (2017). TWEAK/Fn14 Activation Participates in Skin Inflammation. Mediators Inflamm. 2017, 6746870. 10.1155/2017/6746870.

52. Yardman-Frank, J.M., and Fisher, D.E. (2021). Skin Pigmentation and its Control: From Ultraviolet Radiation to Stem Cells. Exp. Dermatol. 30, 560–571. 10.1111/exd.14260.

53. Zenz, R., and Wagner, E.F. (2006). Jun signalling in the epidermis: From developmental defects to psoriasis and skin tumors. Int. J. Biochem. Cell Biol. 38, 1043–1049. 10.1016/j.biocel.2005.11.011.

54. Holleran, W.M., Takagi, Y., and Uchida, Y. (2006). Epidermal sphingolipids: metabolism, function, and roles in skin disorders. FEBS Lett. 580, 5456–5466. 10.1016/j.febslet.2006.08.039.

55. Archambault, M., Yaar, M., and Gilchrest, B.A. (1995). Keratinocytes and fibroblasts in a human skin equivalent model enhance melanocyte survival and melanin synthesis after ultraviolet irradiation. J. Invest. Dermatol. 104, 859–867. 10.1111/1523-1747.ep12607034.

56. Nakazawa, K., Kalassy, M., Sahuc, F., Collombel, C., and Damour, O. (1998). Pigmented human skin equivalent--as a model of the mechanisms of control of cell-cell and cell-matrix interactions. Med. Biol. Eng. Comput. 36, 813–820. 10.1007/BF02518888.

57. Wu, F., Fan, J., He, Y., Xiong, A., Yu, J., Li, Y., Zhang, Y., Zhao, W., Zhou, F., Li, W., et al. (2021). Single-cell profiling of tumor heterogeneity and the microenvironment in advanced non-small cell lung cancer. Nat. Commun. 12, 2540. 10.1038/s41467-021-22801-0.

58. Hao, Y., Hao, S., Andersen-Nissen, E., Mauck, W.M., Zheng, S., Butler, A., Lee, M.J., Wilk, A.J., Darby, C., Zager, M., et al. (2021). Integrated analysis of multimodal single-cell data. Cell 184, 3573–3587.e29. 10.1016/j.cell.2021.04.048.

59. McGinnis, C.S., Murrow, L.M., and Gartner, Z.J. (2019). DoubletFinder: Doublet Detection in Single-Cell RNA Sequencing Data Using Artificial Nearest Neighbors. Cell Syst. 8, 329–337.e4. 10.1016/j.cels.2019.03.003.

60. Germain, P.-L., Lun, A., Meixide, C.G., Macnair, W., and Robinson, M.D. (2022). Doublet identification in single-cell sequencing data using *scDblFinder*. Preprint at F1000Research, 10.12688/f1000research.73600.2

61. Butler, A., Hoffman, P., Smibert, P., Papalexi, E., and Satija, R. (2018). Integrating single-cell transcriptomic data across different conditions, technologies, and species. Nat. Biotechnol. 36, 411–420. 10.1038/nbt.4096.

62. Liu, B., Li, C., Li, Z., Wang, D., Ren, X., and Zhang, Z. (2020). An entropy-based metric for assessing the purity of single cell populations. Nat. Commun. 11, 3155. 10.1038/s41467-020-16904-3.

63. Yu, G., Wang, L.-G., Han, Y., and He, Q.-Y. (2012). clusterProfiler: an R package for comparing biological themes among gene clusters. Omics J. Integr. Biol. 16, 284–287. 10.1089/omi.2011.0118.

64. Carlson, M., Falcon, S., Pages, H., and Li, N. (2019). org. Hs. eg. db: Genome wide annotation for Human. R Package Version 3, 3.

